# High-throughput SWCNT NIR-II screening enables cell and *in vivo* applications

**DOI:** 10.64898/2026.01.27.701967

**Authors:** Atara R. Israel, Zachary Cohen, Nazifa Rahman, Anastasiia Vasylaki, Amelia Ryan, Syeda Rahman, Pratyusha Ghosh, Ryan M. Williams

## Abstract

Diagnostic sensor development and clinical translation lag, in part, due to a lack of rapid, high-throughput screening methodology. Optimization of high-throughput sensor development and deployment will likely help in expediting tools toward the clinic. To address this issue, we optimized high-throughput screening parameters using a near-infrared (NIR-II) plate reader attached to an external probe for *in vivo* testing. We assessed spectroscopy parameters to improve speed and precision in screening a single-walled carbon nanotube (SWCNT)-based optical sensor. To do so, we assessed the appropriate well plate specifications, including laser power, excitation wavelength, exposure time, and focal height parameters for SWCNT-based optical sensor development. We also used the plate reader to screen fluorescent SWCNT which were endocytosed by a macrophage cell line. We then performed NIR probe spectroscopy to assess SWCNT embedded within a methylcellulose hydrogel. Finally, we used the NIR probe to measure SWCNT center wavelength and intensity from live immunocompetent mice. We anticipate that this framework may be broadly applicable to the development of near infrared nanosensors with the potential for more rapid clinical diagnostic translation.

## Introduction

Near infrared (NIR) spectroscopy is a valuable tool for biomedical applications, including noninvasive deep-tissue imaging and non-destructive chemical analysis of biofluid samples^1^. NIR spectroscopy enables rapid, low-cost diagnostic tools without sacrificing selectivity and sensitivity for clinical applications^2^. Alternative clinical diagnostic methods have significant disadvantages in that many are labor intensive, time consuming, and costly. Imaging modalities such as MRI, CT, and PET have notable drawbacks, including low temporal resolution, exposure to ionizing radiation, and incompatibility with point-of-care or bedside diagnostics. The NIR-II region of the electromagnetic spectrum (1000-1700 nm), commonly referred to as the biological tissue transparency window, is especially attractive for supplementary diagnostic purposes through biological imaging. This region of the electromagnetic spectrum exhibits high tissue penetration depth with minimal water and photon absorption, along with minimal tissue autofluorescence and light scattering^3-6^. In most cases, a NIR fluorophore is necessary to transduce information. Applications of NIR-II spectroscopy include quantifying contents of biofluids, measuring oxygen levels in blood, and biological imaging^7^. Some more recent applications include tumor delineation, vascular imaging, photothermal/photodynamic therapy, and drug delivery^8^. Further, NIR spectroscopy is a straightforward method in comparison to other modalities such as Raman spectroscopy, which requires more training and technical expertise^9^.

Many studies have sought to expand the library of NIR fluorescent probes for diagnostic purposes and *in vivo* imaging. NIR-I fluorescent dyes (700 – 1000 nm emission) such as methylene blue and indocyanine green are the only two NIR dyes approved by the FDA. Each has been used in several clinical applications, including breast cancer diagnosis, lymphatic mapping, and visualization of tissue perfusion^10, 11^. Many NIR-I organic dyes, such as rhodamine and squaraine, exhibit photobleaching, rapid degradation, or off-target uptake in vivo^5, 12^. Beyond traditional organic fluorophores, a prominent example of a NIR fluorophore is quantum dots (QDs), which have substantial potential for *in vivo* and *in vitro* applications^12^. Rare-earth-doped nanoparticles (RENPs) are another type of probe that have significant advantages in imaging, including a large Stokes shift, long fluorescence lifetime, and minimal photobleaching, though their cost of production and low materials abundance limit their application^5, 8^. Engineered NIR fluorescent proteins have not been widely adopted due to low quantum yields^5^.

Semiconducting single-walled carbon nanotubes (SWCNTs) hold great promise as optical transducers for biosensors, both *in vitro* and *in vivo*^13^. SWCNTs are a quasi-one-dimensional nanomaterial that exhibit NIR-II fluorescence due to a large band gap in their electronic structure^14, 15^. SWCNT fluorescence emission is photostable, non-photobleaching, and exhibits a large Stokes shift. Further, it is highly sensitive to perturbations in its surrounding electrochemical environment, leading to solvatochromism or changes in brightness in response to external stimuli^16^. SWCNT structures are described as chirality, or (*n,m*) index, defined by diameter and chiral angle^17^. Each (*n,m*) has a unique electronic structure, leading to a distinct emission wavelength. These optical properties have enabled the design of many SWCNT-based optical sensors for a variety of biological analytes, including small molecules^18-25^, macromolecules^26^, proteins^27-30^, lipids^31^, enzymes^32^ and nucleic acids^33, 34^. Furthermore, several SWCNT-based sensors have been used *in vivo* due to the ability of NIR-II fluorescence to penetrate through biological tissues. There are several ways in which SWCNTs can be implemented for *in vivo* use with the potential to realize rapid sensing, enhanced personalized treatment, and robust output^19, 35-38^.

SWCNT-based nanosensor development employs both screening approaches and rational design with known biomolecular recognition elements. The most common example of a screening approach is the corona phase molecular recognition (CoPhMoRe) method. This entails screening a library of organic polymers^39^ or DNA sequences^40^ that can simultaneously solubilize SWCNT and create binding affinity to a specific analyte. Another example is high-throughput screening coupled with machine learning to ascertain spectral disease signatures^41-44^. Both examples require substantial screening of analytes and recognition probes, typically in triplicate; though machine learning-based approaches require many thousands of data points. Rational design of SWCNT-based sensors is also possible, typically using aptamers^27^, antibodies^28-30, 45^, peptides^46, 47^, or other elements with known binding affinity for the desired target. In these studies, high-throughput sensor analysis is required to iterate through sensor design and evaluate standard sensor characteristics.

It is clear that high-throughput instrumentation is necessary to screen and efficiently develop high-performing nanosensors. Such instrumentation is widely available on the market for visible fluorophores, but only emerging for NIR fluorophores. Further, to make use of the unique properties of SWCNT to study disease biology or develop translational sensors, *in vivo* deployment and validation is necessary. While there are examples of visible whole-animal imaging systems, there are few, if any, examples of rapid spectroscopic data collection, largely due to the autofluorescence of tissue in the visible range. In this work, we optimized experimental conditions for plate reader-based high-throughput screening in solution and cells to in vivo used via an integrated workflow framework.

## Methods

### High-throughput screening and in vivo spectroscopy combined instrumentation

High-throughput data collection was obtained via absorbance and fluorescence spectroscopy using the ClaIR NIR-II Microplate Reader with attached point-and-shoot IRina spectral probe (Photon etc.; Montreal, QC) (**Figure 1**). The system contains two diode lasers at excitation wavelengths of 655 nm and 730 nm, with a maximum output in the microplate reader of 3 W in a continuous wave, and 1 W for the spectral probe. The microplate reader can measure NIR-II spectra from 900-1600 nm and absorbance spectra from 500-1600 nm using a halogen white lamp. The distance of the well plate height from the bottom of the laser can be adjusted with a 0.1 mm resolution. The spectral probe contains a spectral resolution of 5 nm. The ClaIR system runs on independent ClaIR software, while the IRina system operates on separate PHySpec software (Photon etc.).

**Figure 1.**
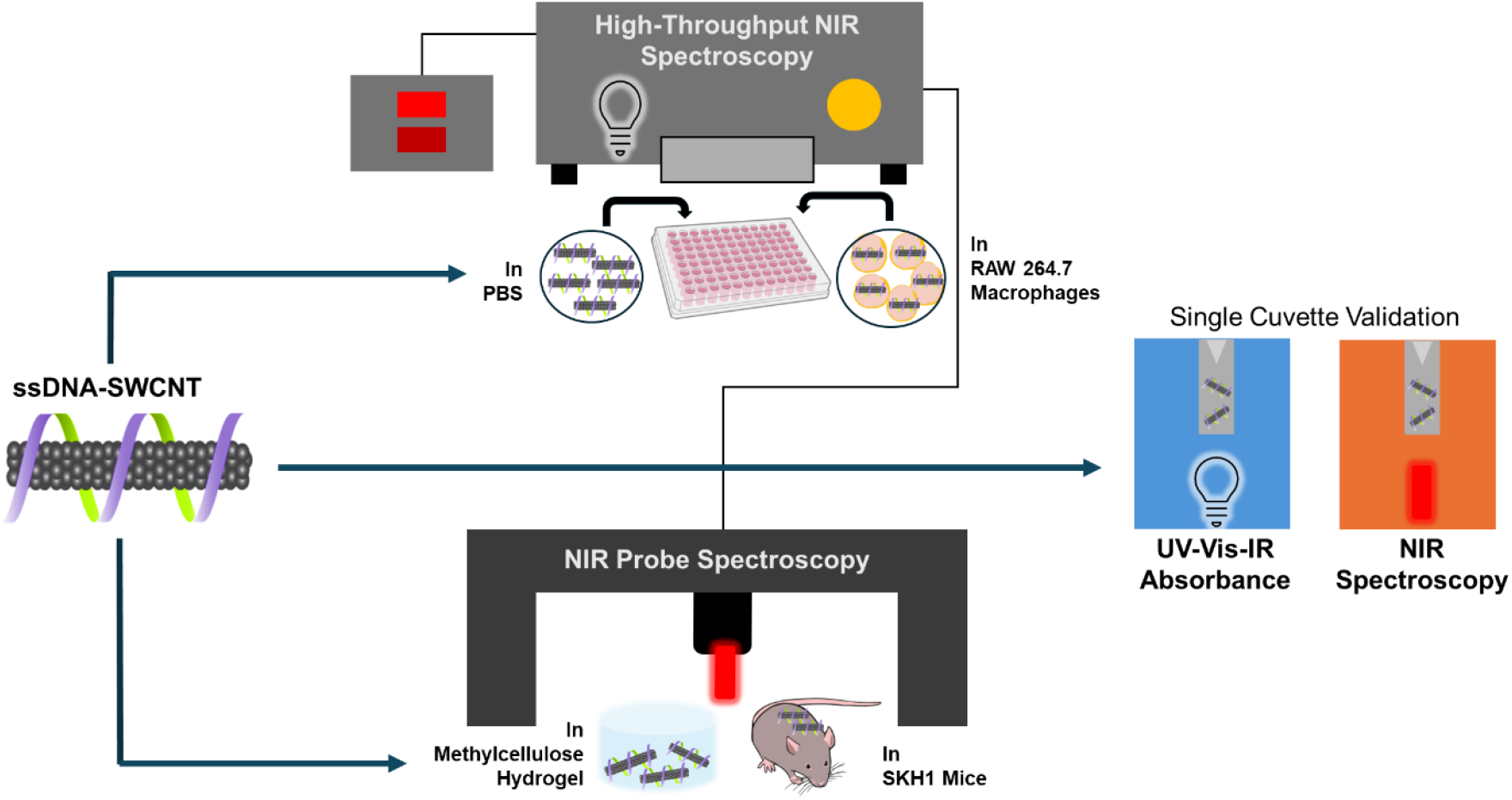
Schematic of fluorescent SWCNT screening. High-throughput plate reader-based NIR spectroscopy was performed for SWCNT in buffer and cells. NIR probe spectroscopy was performed for SWCNT embedded into a hydrogel and in live mice. Spectral measurements were validated with single cuvette instrumentation.

### Validation with single cuvette spectroscopy

To validate plate reader measurements, we compared them to measurements from single-cuvette NIR spectroscopy and a single-cuvette UV-Vis spectrometer. We used a V-730 UV–Vis Spectrometer (Jasco; Easton, MD) for visible absorbance measurements and Nano Spectralyzer (NS) MiniTracer (Applied NanoFluorescence; Houston, TX) single-cuvette NIR absorbance and fluorescence spectrometer. For both instruments, single-use BrandTech UV-Cuvettes (Fisher Scientifc; Hampton NH), were used, with 1x PBS as a reference. The absorbance measurements taken on the UV-Vis spectrometer were taken with a scan speed of 400 nm/min. The absorbance measurements on the NIR spectrometer were taken with an integration time of 40 counts/s averaged over 3 measurements, while the fluorescence measurements were taken with a 1000 count/s integration time averaged over 3 measurements.

### Fluorescent SWCNT formulation

HiPCo prepared SWCNT (NanoIntegris Technologies; Boisbriand, QC) were dispersed in a 1% sodium deoxycholate solution (DOC; Fisher Scientifc; Hampton NH) using probe tip sonication^48^ with a 1/8” microtip probe (Sonics; Newtown, CT), followed by ultracentrifugation at 58,000 X *g* for 1 hour using an Optima Max-XP Ultracentrifuge (Beckman Coulter; Brea, CA). The top 75% of the supernatant was collected and stored at 4°C until ready for use. For ssDNA (Integrated DNA Technologies; Coralville, IA) instead of DOC suspension, an additional filtration step to remove free DNA was done using a 100 kDa Amicon centrifugal filter (Sigma-Aldrich; St. Louis, MO) for 15 min at 14,000 X *g* and resuspended in 100 µL 1x PBS.

### Screening commercial well plates for high-throughput measurements

To find the optimal transparent plate for near-infrared sample measurements, five were tested for well-to-well variation in spectral acquisition. Tested plates included the Corning UV transparent half area 96-well plate (3679; Fisher Scientific; Hampton NH), Brandtech 96-well plate (781611; Fisher Scientific; Hampton NH), Corning 384-well plate (3640; Fisher Scientific; Hampton NH), Brandtech 384-well plate (781627; Fisher Scientific; Hampton NH), and Corning 1536 well plate (3831; Fisher Scientific; Hampton NH). Three plates of each type were filled with 0.5 mg/L DOC-SWCNT: 7.5 µL in the 1536-well plate, 100 µL in Corning 96-well, 384 well, and BrandTech 384-well plates and 150 µL in BrandTech 96-well plate, into a total of 16 wells per plate. 96 well plates skipped every 3 columns and 1 row, whereas 384 well plates skipped every 6 columns and 3 rows. This was to ensure that there was no variation in the signal acquisition relative to well position on the plate. To reduce variability, we also evaluated the impact of single channel vs multi-channel pipette in plate preparation, along with cleaning the bottom of the plate with a KimWipe (Fisher Scientific; Hampton NH) prior to acquisition. Following plate preparation, we acquired fluorescence spectra with 200 mW laser power with an exposure time of 1000 ms with full emission spectra obtained between 900-1400 nm.

### Optimizing parameters for optical spectroscopy screening

We next sought to optimize optical SWCNT characterization for high-throughput analysis by adjusting laser power, exposure time, gain, and binning. To test these parameters, we used the Corning half area UV transparent 96 well plate for all experiments except for those involving cell culture. We captured absorbance and fluorescence spectra of a 1% DOC-SWCNT dispersion. We modified exposure time between 1 and 1000 ms for both visible and NIR absorbance. Then, we modified laser power from 200-1750 mW, and exposure time from 5-500 ms for fluorescence measurements.

### High-throughput spectroscopy of SWCNT endocytosed by macrophages

Next, we sought to adapt this methodology for high-throughput SWCNT screening in living cells. We investigated whether modification of instrument focal height for surface-attached cell cultures was necessary, as the standard laser-to-plate distance is 500 μm. To determine the optimal focal height, we incubated RAW 264.7 murine macrophages (ATCC; Rockville, MD) at 80% confluency with 10 mg/L ssDNA (GT)_15_-SWCNT for 30 minutes in a tissue-culture treated transparent 96 well plate (Fisher Scientific; Hampton NH), followed by two washes with 1x PBS and addition of fresh media^43^. Fluorescence measurements were acquired using a laser power of 1750 mW and exposure time of 500 ms, with the z-height adjusted between 0-1 mm to determine the influence of distance between the bottom of the plate and the laser on the resulting spectra.

### Assessment of hydrogel-encapsulated SWCNT using a spectroscopic probe attachment

One potential use of SWCNT *in vivo* is encapsulated within a hydrogel. To optimize use of the spectral probe, we encapsulated 1 mg/L (GT)_15_-SWCNT within methacrylated methylcellulose hydrogels^19^. We cast gels into a black Brandtech 96 well plate with a transparent bottom prior to gelation at room temperature for 30 minutes. Six out of 12 wells were topped with 100 µL 1x PBS to mimic analyte addition. We adjusted exposure time from 250-1000 ms, at 1000 mW, and power from 250-1000 mW, at 1000 ms to optimize data collection. A single acquisition of the well was first taken and used as a background for subtraction in data analysis.

### Rapid spectral analysis in live mice

We next evaluated optimal fluorescence spectral acquisition parameters in live mice. All experiments were approved by the Institutional Animal Care and Use Committee of The City College of New York. We subcutaneously injected 2 mg/L (GT)_15_-SWCNT into the flank of three hairless, immunocompetent, SKH1-*Hr*^*hr*^ female mice (4-6 weeks old)^36^. Mice were anesthetized with vaporized isoflurane for spectra acquisition. Spectral measurements were acquired immediately after injection, 1 hour, 4 hours, and 24 hours post-injection using a laser power of 1000 mW with an exposure time of 1000 ms. A single acquisition was taken of the area being measured prior to injection and used as a background. Further data processing was performed in OriginPro.

### Data collection, analysis, and statistics

All data was collected in triplicate, unless otherwise noted, with the mean ± standard deviation reported. Peak fitting analysis was performed on the fluorescence spectra using a custom MATLAB code (available upon request) to fit the peaks to a pseudo-Voigt profile and obtain center wavelength and maximum intensity values for each chirality. Fits with R^2^ > 0.99 were considered successful. Standard square error (SSE) and root mean square error (RMSE) were also calculated for each fit.

## Results and Discussion

Using a high-throughput NIR-II microplate reader, we obtained representative absorbance and fluorescence spectra of DOC-SWCNT (Figures 2A-C). Absorbance spectra were captured over the visible and IR regions, from 500-1600 nm. Fluorescence emission spectra were acquired with both 655 and 730 nm excitation wavelengths between 900-1700 nm. The representative absorbance spectra show signature peaks at 1150 nm and 1250 nm, while fluorescence spectra display the characteristics peaks of SWCNT for the (7,6), (7,5) (655 nm) and (9,4) (730 nm) chiralities at 1040 nm, 1137 nm, and 1129 nm, respectively.

**Figure 2:**
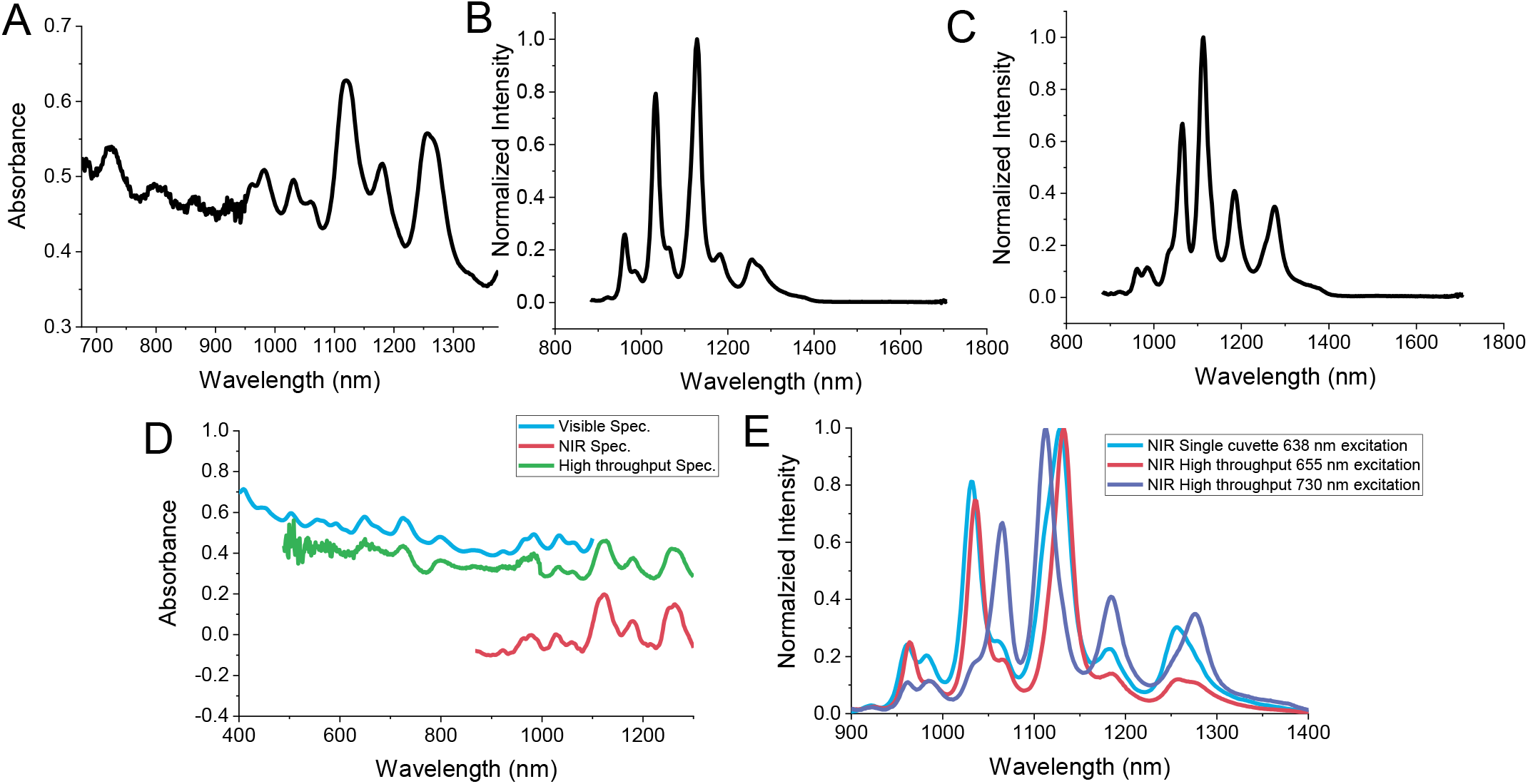
Representative spectra of SWCNT in a high-throughput NIR plate reader. A) Single representative absorbance spectra of 10 mg/L DOC-SWCNT using an exposure time of 1.75 ms Vis, 75 ms NIR. B) Single representative fluorescence spectra of 10 mg/L DOC-SWCNT, excited by a 655 nm laser source. Fluorescence parameters: laser power 200 mW, exposure time 500 ms C) Single representative fluorescence spectra of 10 mg/L DOC-SWCNT, excited by a 730 nm laser source. Fluorescence parameters: laser power 200 mW, exposure time 500 ms. D) Comparison of representative absorbance spectra taken on the visible spectrometer, NIR single cuvette spectrometer, and high-throughput spectrometer. E) Comparison of representative fluorescence spectra taken on the single cuvette NIR spectrometer at an excitation of 638 nm and high-throughput spectrometer at excitation wavelengths of 655 and 730 nm.

Absorbance acquisition using the single-cuvette visible spectrometer provides spectral measurements over the range of 300-1100 nm, while the single-cuvette NIR-II spectrometer collects absorbance between 900-1600 nm. (Figure 2D). While both single-cuvette instruments collect absorbance of samples over their respective ranges, we obtained continuous absorbance measurements from the plate reader over both the visible and near-infrared regions. Comparing the IR absorbance collected using the single-cuvette vs microplate reader, a higher signal is obtained with the latter. However, visible absorbance collected with the single-cuvette visible spectrometer yields a slightly higher signal than the plate reader.

Fluorescence spectra acquired on the single-cuvette spectrometer in comparison to the high-throughput spectrometer has some differences; namely the excitation source of the instruments differ by 17 nm (single cuvette excitation = 638 nm, plate reader excitation = 655 nm). Because of this, the characteristic peaks of SWCNT may be resolved at slightly different wavelengths (Figure 2E). The power of the laser source is also a point of variance between the instruments: while the single cuvette spectrometer has a fixed power of 50 mW, the plate reader spectrometer allows laser power to be adjusted from 200-1750 mW.

### Determining use of appropriate commercial well plates

While most well plates are manufactured for use with visible absorbance and fluorescence plate readers, we sought to determine which of several commonly-used options were most compatible for NIR-II data collection. We tested five different plates: Corning half area 96-well (UV transparent), Brandtech 96-well, Corning 384-well, Brandtech 384-well, and Corning 1536 well plate. Comparing the variance in center wavelength and intensity values among the plates tested for the (7,5) and (8,6) chiralities shows that the Corning 96-well half area plates resulted in the lowest maximum variance (0.098nm) compared to the other plates used (Figure 3). Additionally, we loaded 16 wells of 3 Corning 96-well plates with both a single-channel pipette and multi-channel pipette to compare the impact on the fluorescence signal for each sample. Based on the variance in peak intensity and center wavelength for the (7,5) and (8,6) chiralities, the multi-channel pipette produced lower maximum variance for both chiralities (0.006 and 0.039 nm; 1.07 and 1.04 ADU). Another potential point of variance in plate preparation is the bottom of the well plate collecting ambient dust, which may interfere with signal acquisition. However, we found no difference in Brandtech 96-well plate fluorescence spectra before or after gentle wiping of the bottom surface. Overall, the variance of center wavelength shifts and intensities points to the Corning half area 96-well plates to be the most compatible with the NIR-II plate reader (Figure 3). The plate being rated for UV use is also beneficial for absorbance acquisition using the microplate reader, along with fluorescence measurements. The Corning 96-well plate has also been used in previous experiments of similar design with different instruments, demonstrating its compatibility and utility for nanotube-based fluorescence detection^20, 28, 29^.

**Figure 3.**
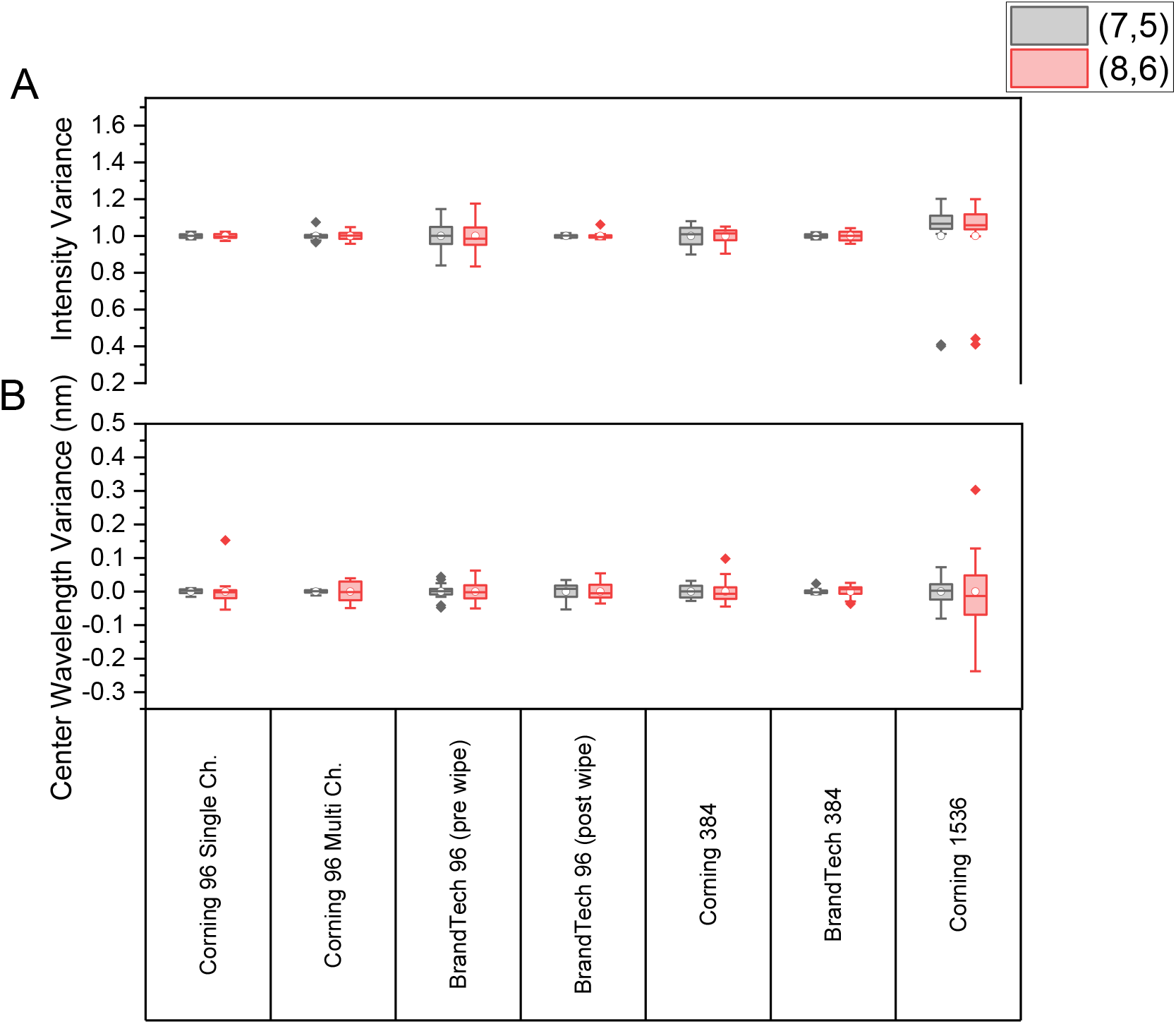
Variance in sample measurements from various well plates. A) Intensity variance of the (7,5) and (8,6) chiralities of 0.5 mg/L 1% DOC-SWCNT. B) Variance in the center wavelength of the (7,5) and (8,6) peaks of 0.5 mg/L 1% DOC-SWCNT.

### Optimal sensor parameters for high-throughput screening

We next tested a range of light and laser exposure times on 1 mg/L DOC-SWCNT to evaluate their impact on the resulting absorbance and fluorescence. For absorbance spectra, exposure times between 1-1000 ms were tested for both visible and IR absorbance. As exposure time increased between 10-100 ms, the absorbance peaks became smoother and brighter, as expected (Figure 4A). The maximum exposure time for absorbance measurements on the high-throughput plate reader is 1000 ms. Very low exposure times (∼10 ms) need to be used to get a proper signal of SWCNT in the visible region (Figure 4B), otherwise the characteristic peaks become oversaturated. Increasing IR absorbance exposure time to 100 ms does improve spectral acquisition (Figure 4C), displaying a spectrum with smooth, defined peaks in the IR range. While exposure time selection will vary based on the sample concentration, it is ideal to use lower times (1-10 ms) to obtain readable spectra in both the visible region and higher times (50-250 ms) in the IR region.

**Figure 4.**
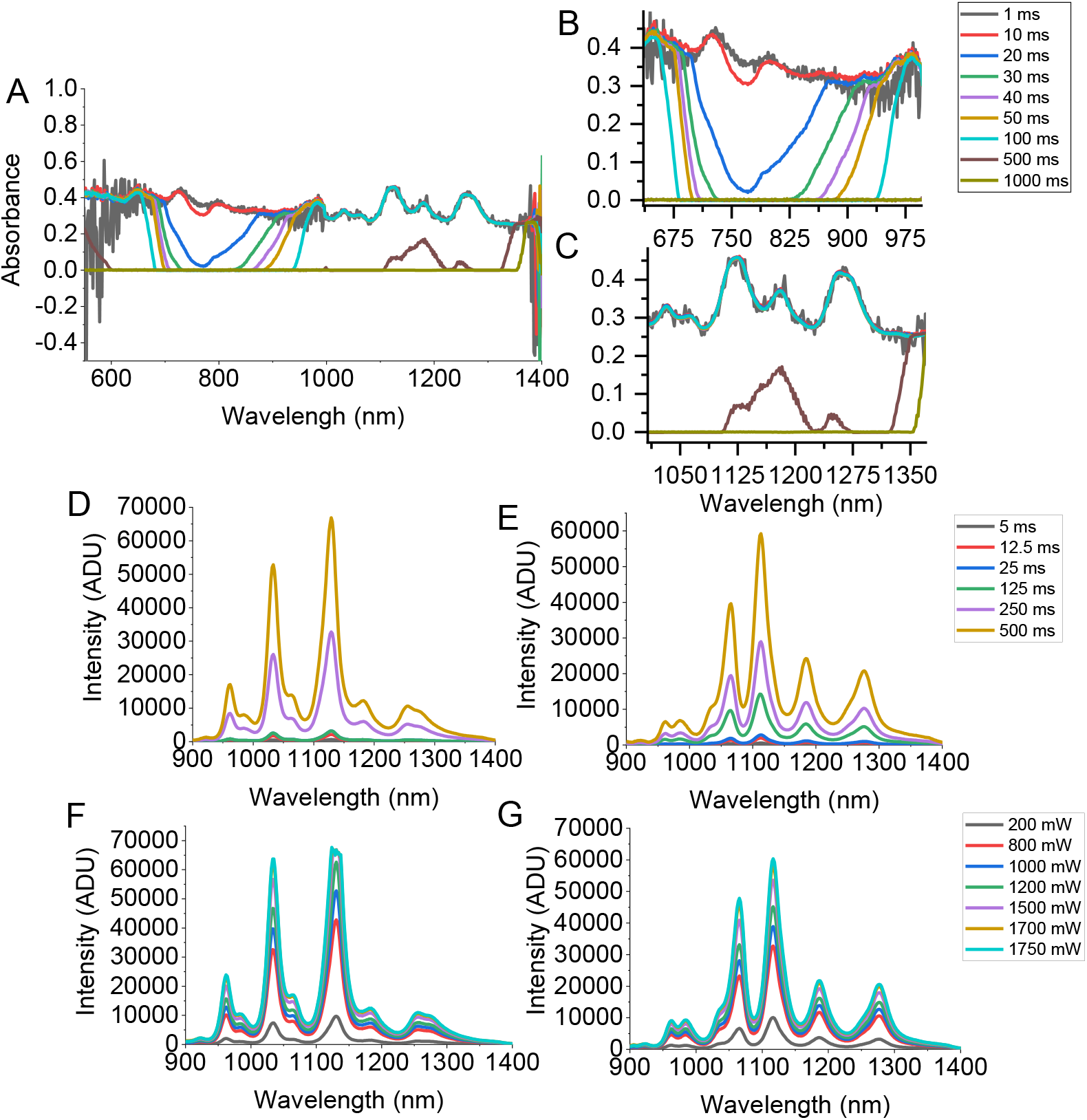
Optimizing spectral acquisition parameters. A) Full spectra of absorbance measurements taken on the high-throughput plate reader at exposure times ranging from 1-1000 ms in both the visible and near infrared regions. B) Close-up of visible absorbance region of SWCNTs. C) Close-up of IR absorbance region of SWCNTs. D) Exposure time variation of SWCNT fluorescence spectra at 200 mW laser power for excitation at 655 nm. E) Exposure time variation of SWCNT fluorescence spectra at 200 mW laser power for excitation at 730 nm. F) Laser power variation of SWCNT fluorescence spectra at 1000 ms exposure time at an excitation of 655 nm. G) Laser power variation of SWCNT fluorescence spectra at 1000 ms exposure time at an excitation of 730 nm.

For fluorescence spectroscopy, exposure times ranging from 5-500 ms were tested using DOC-SWCNT at 10 mg/L for excitation wavelengths 655 nm and 730 nm, with laser power remaining constant at 200 mW. As expected, increasing exposure time to the sample increases the intensity of the SWCNT fluorescence signal for each of the 655 and 730 nm lasers (Figures 4D and E). Upon pseudo-Voigt peak-fitting analysis of the fluorescence spectra, we found that higher exposure times yielded lower standard deviation of the center wavelength and low root mean square error (RMSE) and sum of squared error (SSE) values across samples at different chiralities, while the standard deviation of intensity values saw an increase with higher exposure times (Supplementary Figure 1). This indicates consistent measurements among samples for spectra obtained at higher exposure times.

We then examined the effect of laser power on spectral acquisition using 1 mg/L DOC-SWCNT. With a fixed exposure time of 500 ms, the laser power for both 655 nm and 730 nm excitation wavelengths were adjusted between 200-1750 mW. As expected, SWCNT fluorescence intensity increased with increasing laser power (Figure 4F and G). At around 1500 mW and above for the 655 nm laser, the (7,6) peak became overexposed, saturating the detector. The standard deviation of the peak center wavelength for most chiralities analyzed showed minimal differences as power increased. The standard deviation for maximum intensity values, however, did increase with increasing laser power from 45 up to 2400 ADU. Lower RMSE and SSE values were also observed with increased laser power (Supplementary Figure 2). Nevertheless, using higher laser power does produce more defined peaks. Based on these results, we found that it is best to use laser powers above 1500 mW with our samples to obtain brighter signals. However, subsequent adjustment of the exposure time based on sample concentration is needed to avoid overexposure.

We also evaluated the effect of 2×1 data binning during spectral acquisition when adjusting the laser power. Data binning is a preprocessing method that groups a range of numbers into bins to condense the data in an effort to increase the signal to noise ratio. 2×1 binning was applied to the spectrometers during acquisition, summing the pixel values into pairs and reducing the resolution of the output signal by a factor of 2. For the (8,3), (7,5), (9,5+10,3), (10,2), (9,4), and (8,6) peaks, the un-binned data points had lower standard square error (SSE) and root mean square error (RMSE) values compared to binned data points (Supplementary Figure 6). This may be explained by the abundance of certain chiral structures in the SWCNT mix; the more abundant species of nanotubes, such as the (7,5) and (9,4) chiralities, produce higher peaks and therefore more data points are obtained for those peaks. Less prominent peaks, such as those from the (9,5+10,3), and (8,7) chiralities, produce less intense fluorescence, thus binning the data points reduces the information obtained from those peaks. While enabling binning produces smoother fitted peaks, it does reduce the resolution of the data and should therefore only be applied to spectra that are initially noisy. With binning enabled during spectral acquisition, the number of data points collected over the full emission spectrum is half of that acquired without binning (Supplementary Figure 3A). Therefore, when data is obtained across the NIR-II spectrum, we found that it is best to do so without binning and apply it as necessary for analysis and if needed after data collection.

The NIR-II plate reader system integrates the ability to acquire spectra in high gain mode, which may be used for signals that are dim and require extra amplification. We tested the use of the high gain mode with low laser powers (200 mW). However, we found that the high gain mode produced a noisy signal with undefined peaks (Supplementary Figure 3A). High gain mode can be useful to amplify signals from samples that may be too dim or noisy to read initially, or when high laser powers and exposure times cannot be used if they negatively affect the sample.

### Screening fluorescence of SWCNT endocytosed by macrophages

We next sought to optimize spectral acquisition of (GT)_15_ ssDNA-SWCNT endocytosed by RAW 264.7 macrophages. Prior work has used imaging methodologies coupled with machine learning to phenotype macrophages with (GT)_15_-SWCNT^43^, while other studies have evaluated surface binding of SWCNT to cells in a high-throughput manner^49^. For experiments involving cell culture, measuring SWCNT uptake and spectra may require nanotube excitation to take place at a closer distance to the sample. The default laser-to-plate distance of the plate reader is 500 µm. We investigated whether a smaller distance between the bottom of the plate and the laser would increase the fluorescence intensity of SWCNTs endocytosed by macrophages, enhancing signal. We therefore modified the distance as allowed by the instrument (0-1 mm), finding some changes in the spectral signal of SWCNT inside of cells. The signal was slightly dimmer at 1 mm and was only marginally increased at 750 µm and 10 µm. These differences are negligible and likely represent experimental variance, as no clear trend was observed (Figure 5). This experiment does, however, demonstrate that high-throughput screening of SWCNT inside of cells is possible, potentially allowing ensemble measurements from a single well of cells. This could obviate time-consuming single field-of-view screening, potentiating high-throughput drug screening.

**Figure 5.**
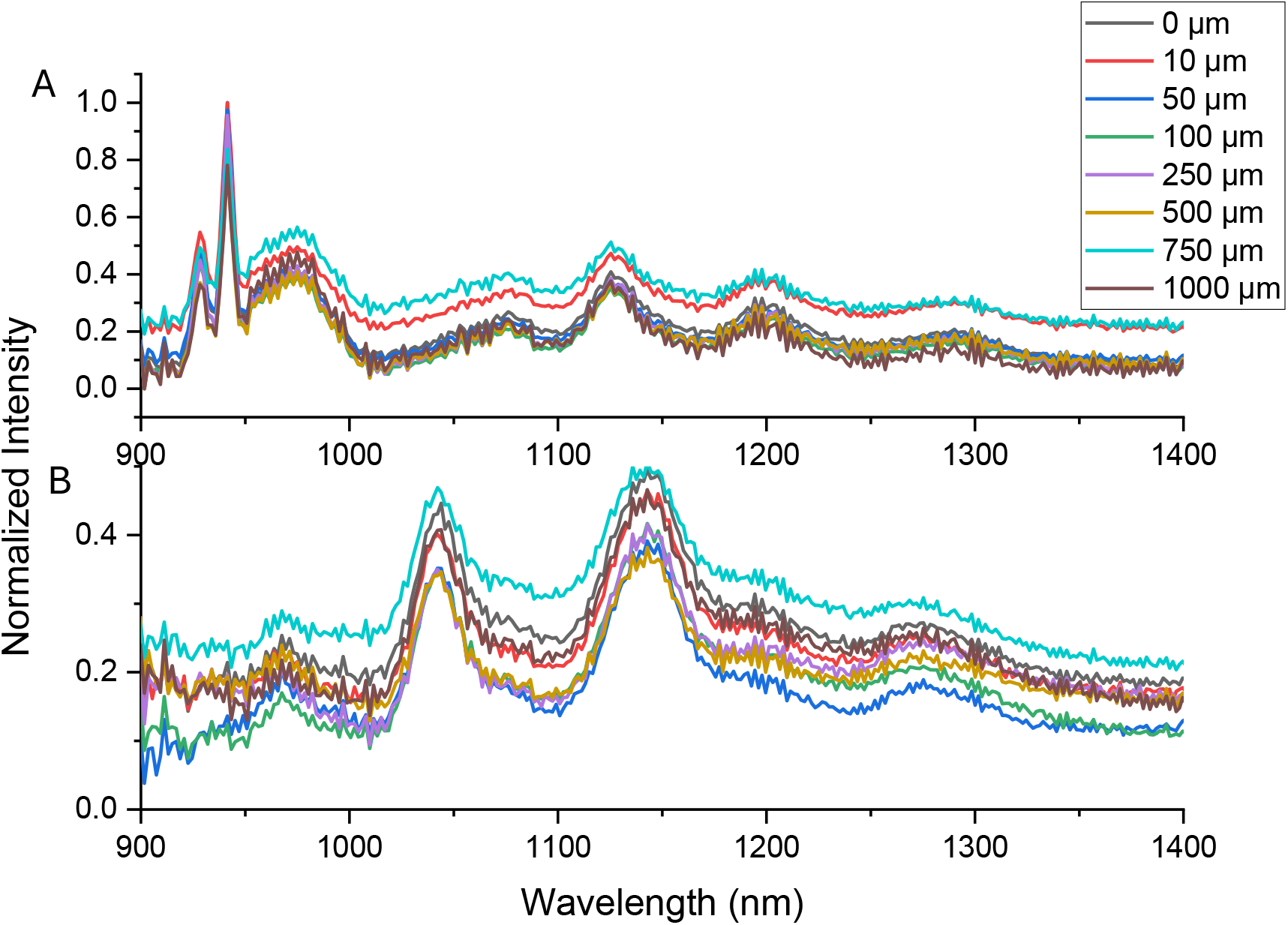
Fluorescence spectra of SWCNTs uptaken by macrophages. Focal height adjustments were made between 0-1 mm at excitation wavelengths of A) 730 nm and B) 655 nm.

### Assessment of hydrogel-encapsulated SWCNT

After sensor lead development in solution or in cells, pre-clinical development would often proceed to live animal validation. To do so, the plate reader is designed to attach to a flexible probe for rapid animal point-and-shoot spectroscopy. To immobilize SWCNT, twelve wells were cast with methylcellulose hydrogels containing 1 mg/L of (GT)_15_-SWCNTs as in our prior work^19^ (Figures 6A and B). We obtained fluorescence spectra with the NIR-II spectral probe with a limited laser power at a maximum of 1000 mW, as we noticed some gel melting above that power, evidenced by gel opacity and inhomogeneity. Observations of the resulting fluorescence spectra reveal that using low laser power and short exposure times produce measurable signal after processing. Peak intensity values with excitation powers of 200 mW and 1000 mW increased by 2.7-fold for 730 nm excitation, and by 4.3-fold for 655 nm excitation. As expected, higher laser powers and longer exposure times produced smooth, narrow peaks (Figures 6C-F). Decreased RMSE and SSE values were observed with increasing exposure time and laser power. The standard deviation of peak center wavelength for all chiralities analyzed decreased with increasing laser power and exposure time. However, standard deviation of peak intensity values increased with laser power and exposure time across chiralities analyzed (Supplementary Figures 4 and 5).

We also investigated the influence of standard overhead fluorescent lighting. Interestingly, we found no substantial differences in peak intensity and center wavelength (Supplementary Figure 3B). This is despite the standard best practice of turning overhead lighting off for such measurements. Automatic binning and high gain modes for signal acquisition were also tested using the spectral probe. High gain mode did not provide any measurable signal (Supplementary Figure 3C). Binning during spectral acquisition slightly lowered the intensity of the signal, but as seen in the plate reader studies, the amount of data points collected with binning enabled is half of what is collected with binning disabled. Binning could be beneficial for samples that exhibit photobleaching or blinking, though resolution of the signal is diminished^50, 51^. High gain mode could be useful for samples that are either very dilute or have dim fluorescence^52^.

**Figure 6.**
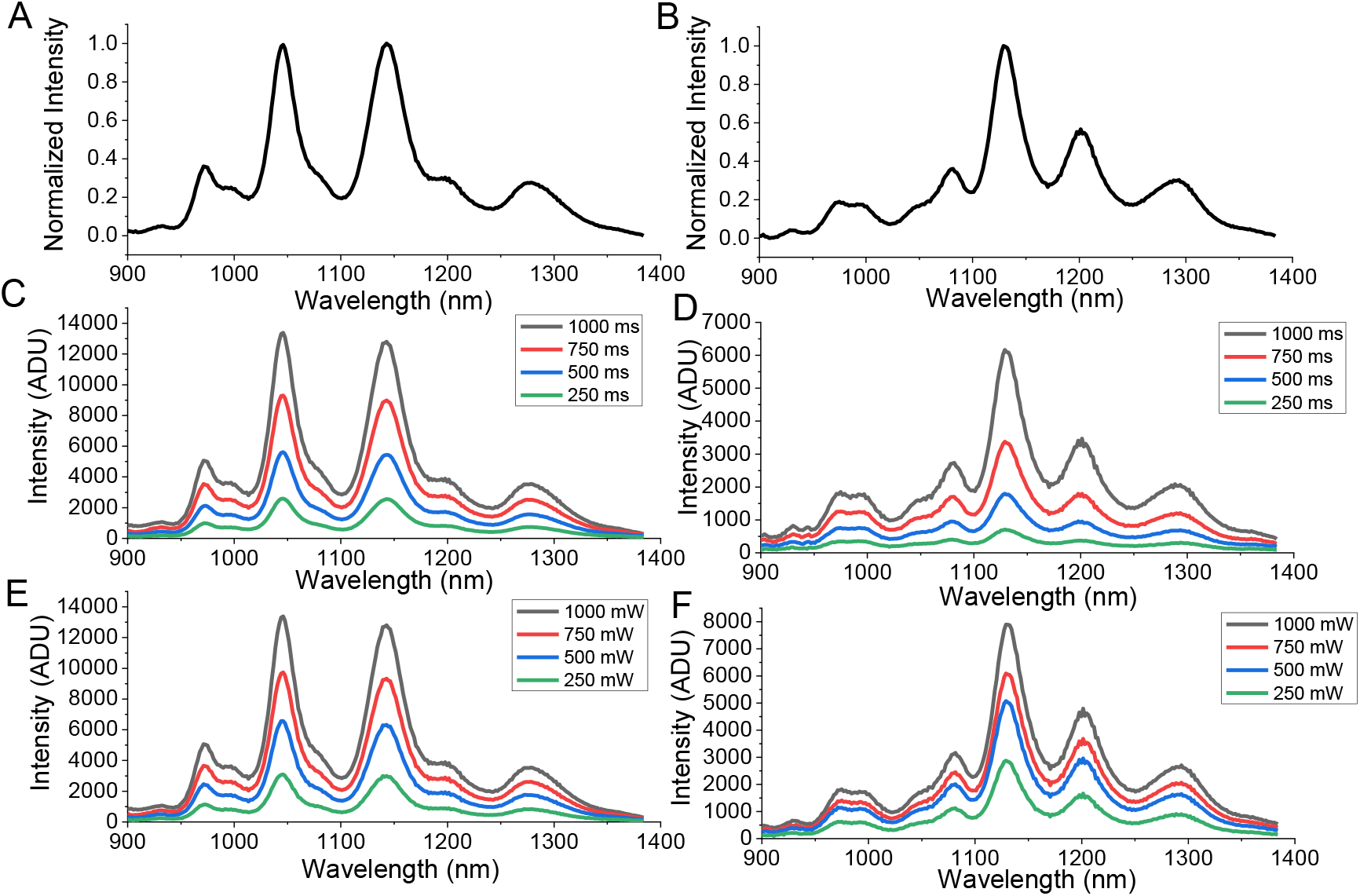
Optimizing spectral probe parameters in a SWCNT-embedded hydrogel. A) Representative spectra of (GT)_15_-SWCNT encapsulated in methylcellulose hydrogels at an excitation of 655 nm B) Representative spectra of (GT)_15_-SWCNT encapsulated in methylcellulose hydrogels at an excitation of 730 nm. C) Exposure time variation of SWCNT fluorescence spectra at 1000 mW laser power for excitation at 655 nm. D) Exposure time variation of SWCNT fluorescence spectra at 1000 mW laser power for excitation at 730 nm. E) Laser power variation of SWCNT fluorescence spectra at 1000 ms exposure time at an excitation of 655 nm. F) Laser power variation of SWCNT fluorescence spectra at 1000 ms exposure time at an excitation of 730 nm.

### Nanosensor detection in live mice

Acquisition of *in vivo* measurements was done by anesthetizing 4-6 week old hairless SKH1-*Hr*^*hr*^ mice and injecting them subcutaneously with 200 µL of 2 mg/L (GT)_15_-SWCNT. Spectral measuements at excitation wavelengths of 655 and 730 nm were taken immediately post injection, 1 hour, 4 hours, and 24 hours post at the site of injection and surrounding area using the spectral probe attachment. We found stable signature SWCNT fluorescence signal after 24 hours at the site of injection (Figure 7A). Peak evaluation of three (*n,m*) ((7,6), (7,5), and (9,4)) also demonstrated a stable fluorescence signal over the timeframe captured (Figures 7B-G).This confirms prior reports that SWCNT spectra remain localized and stable at least over a short time period in live mice^36, 37, 53^. It also validates translation of a lead sensor to testing in a pre-clinical model.

**Figure 7.**
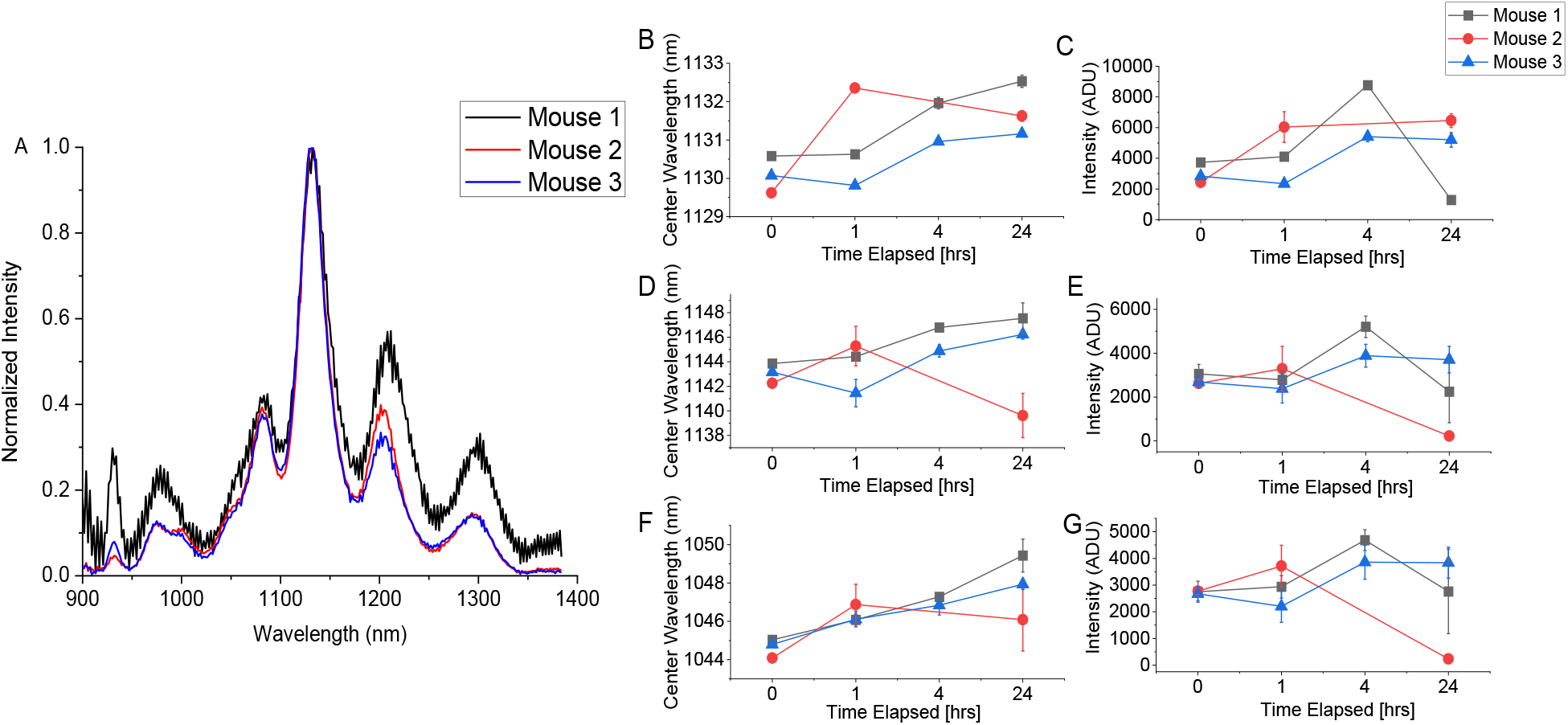
Spectral acquisition *in vivo*. A) Fluorescence spectra at an excitation of 730 nm 24 hours post-SQ injection of (GT)_15_-SWCNT in the flank of mice, average of triplicate measurements per mouse. B-F) Average (n=3) trace of center wavelength and intensity values, respectively, for the B) and C) (9,4) chirality; D) and E) (7,6) chirality; E) and F) (7,5) chirality taken immediately post injection, 1 hour, 4 hours, and 24 hours post injection. Error bars represent ± standard deviation.

## Conclusions

In this work, we optimized high-throughput screening of SWCNT NIR-II sensors in a microplate reader in solution and after endocytosis by macrophages. We then validated spectral acquisition from these sensors after immobilization in a hydrogel and subcutaneous injection into mice. These tools portend the possibility of rapid diagnostic device development, both *in vitro* and *in vivo*. We did so with a model NIR fluorophore, single-walled carbon nanotubes that have stable, bright fluorescence in solution, in cells, and in mice. However, additional optimization may be necessary for other SWCNT formulations, other NIR fluorophores, or solutions. We optimized and validated an expedited framework workflow for pre-clinical lead development and testing. This has the potential to enhance diagnostic probe development and pre-clinical tool expansion.

## Acknowledgements

The authors wish to acknowledge all members of the Williams Lab for discussion and feedback and members of the S. Nicoll Lab at CCNY for assisting with hydrogel preparation. The authors also thank Wendy Chung and Photon etc. for reading and helpful improvements of this manuscript. This work was supported by NIH R35GM142833 and the SUNY Empire Innovation Program (Award #250010) (R. Williams). A. Israel and A. Ryan were supported by a G-RISE Ph.D. traineeship from the National Institutes of Health (T32GM136499).

