## Supplementary Figures for "High-throughput SWCNT NIR-II screening enables cell and *in vivo* applications"

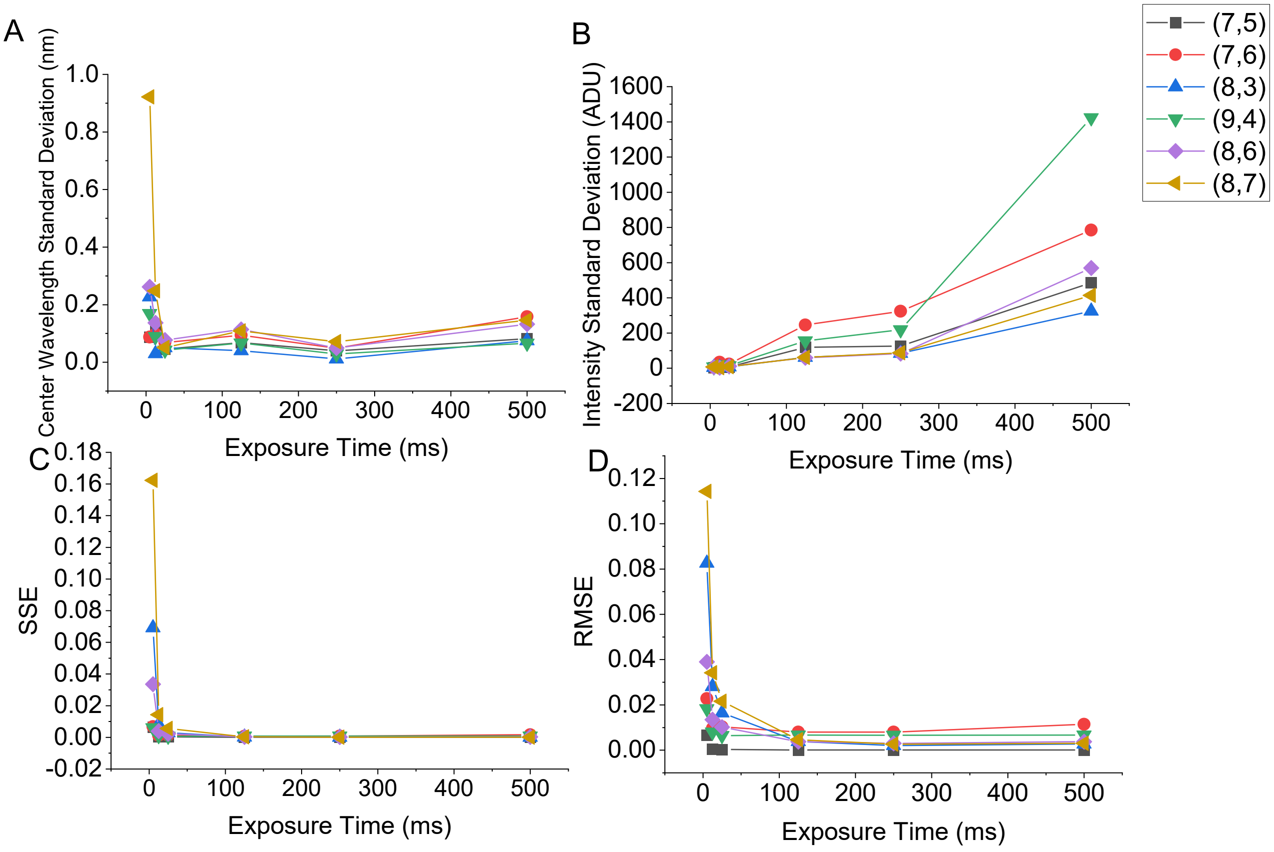


**Supplementary Figure 1**. **Statistical analysis of variations in exposure time of the high throughput system.** A) Standard deviation in center wavelength of each chirality. B) Standard deviation in intensity of each chirality. C) Standard square error (SSE) of each chirality. D) Root mean square error (RMSE) of each chirality. Laser power of 200 mW was used across all time adjustments.

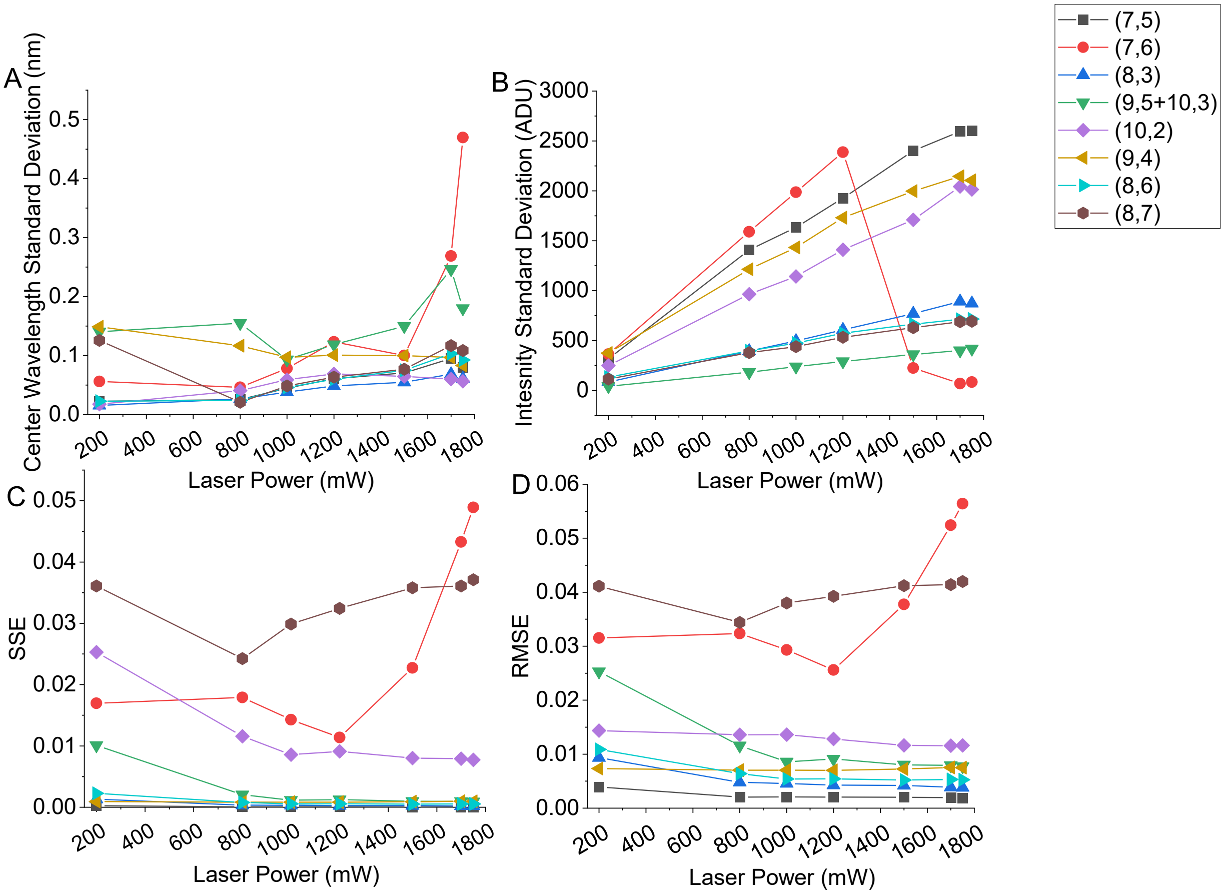


**Supplementary Figure 2. Statistical analysis of variations in laser power of the high throughput system.** A) Standard deviation in center wavelength of each chirality. B) Standard deviation in intensity of each chirality. C). standard square error (SSE) of each chirality. D) Root mean square error (RMSE) of each chirality. Exposure time of 1000 ms was used for all power adjustments.

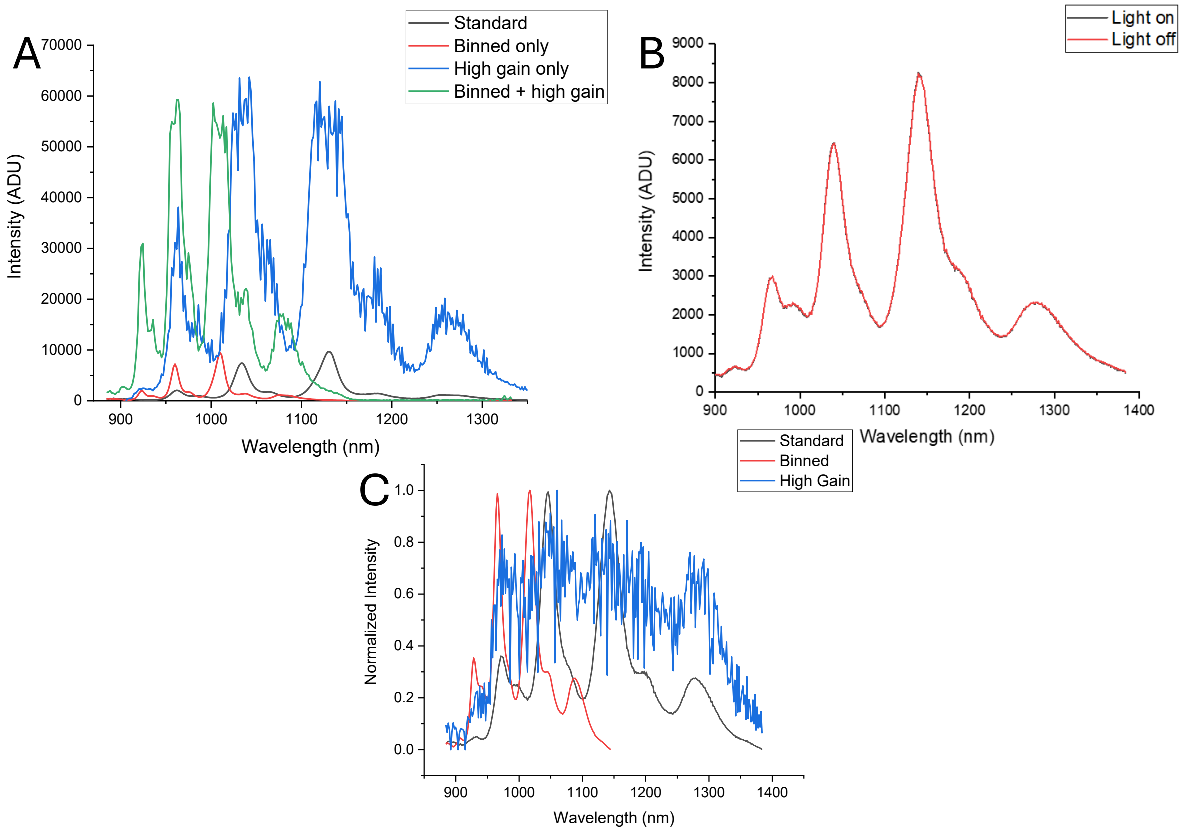


**Supplementary Figure 3. Comparison of alternate conditions on resulting SWCNT fluorescence.** A) Comparison of the resulting fluorescence spectra obtained from the high throughput plate reader using standard parameters (no gain, no bin, 200 mW, 1000 ms), with binning, high gain, and both. B) SWCNT-hydrogel fluorescence spectra taken with lights on and off in the room using the spectral probe. C) Comparison of the resulting fluorescence spectra obtained from the spectral probe using standard parameters (no gain, no bin, 1000 mW, 1000 ms), with binning and high gain mode.

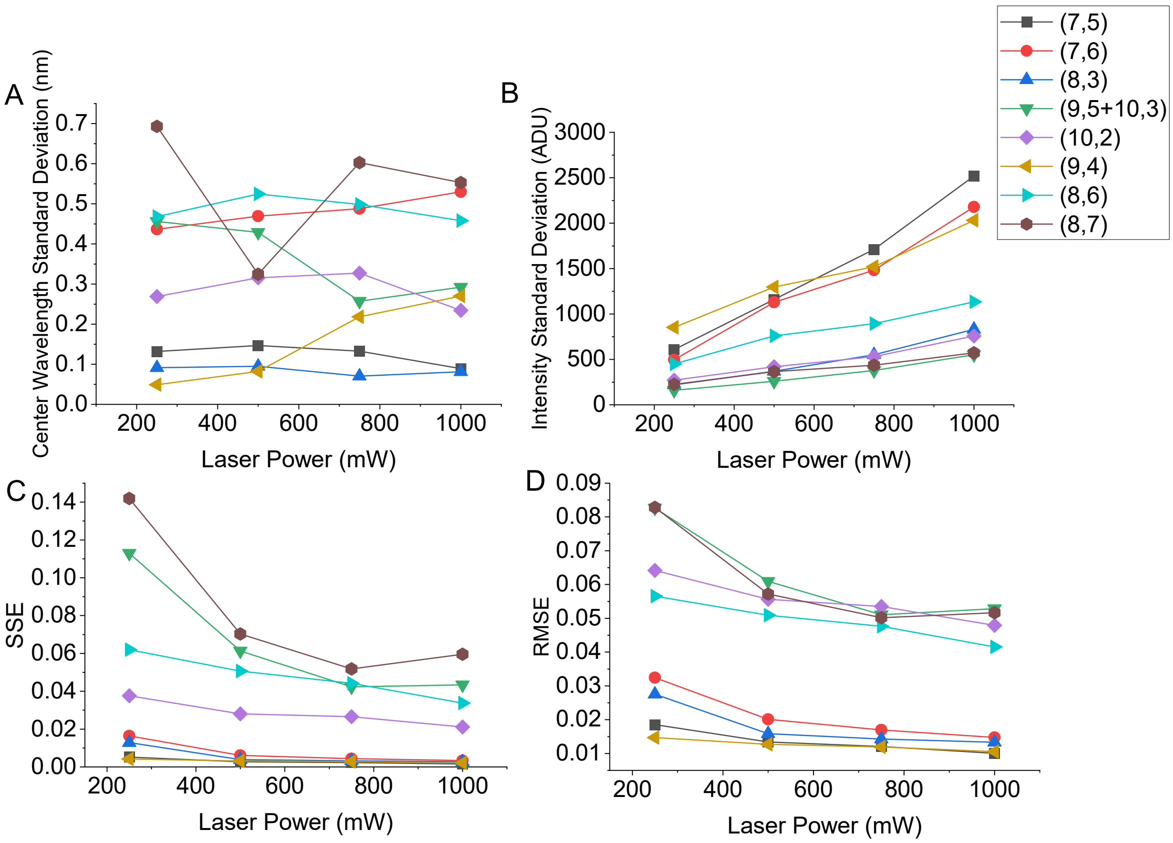


**Supplementary Figure 4**. **Statistical analysis of variations in laser power of the spectral probe system.** A) Standard deviation in center wavelength of each chirality. B) Standard deviation in intensity of each chirality. C). standard square error (SSE) of each chirality. D) Root mean square error (RMSE) of each chirality. Exposure time of 1000 ms was used for all power adjustments.

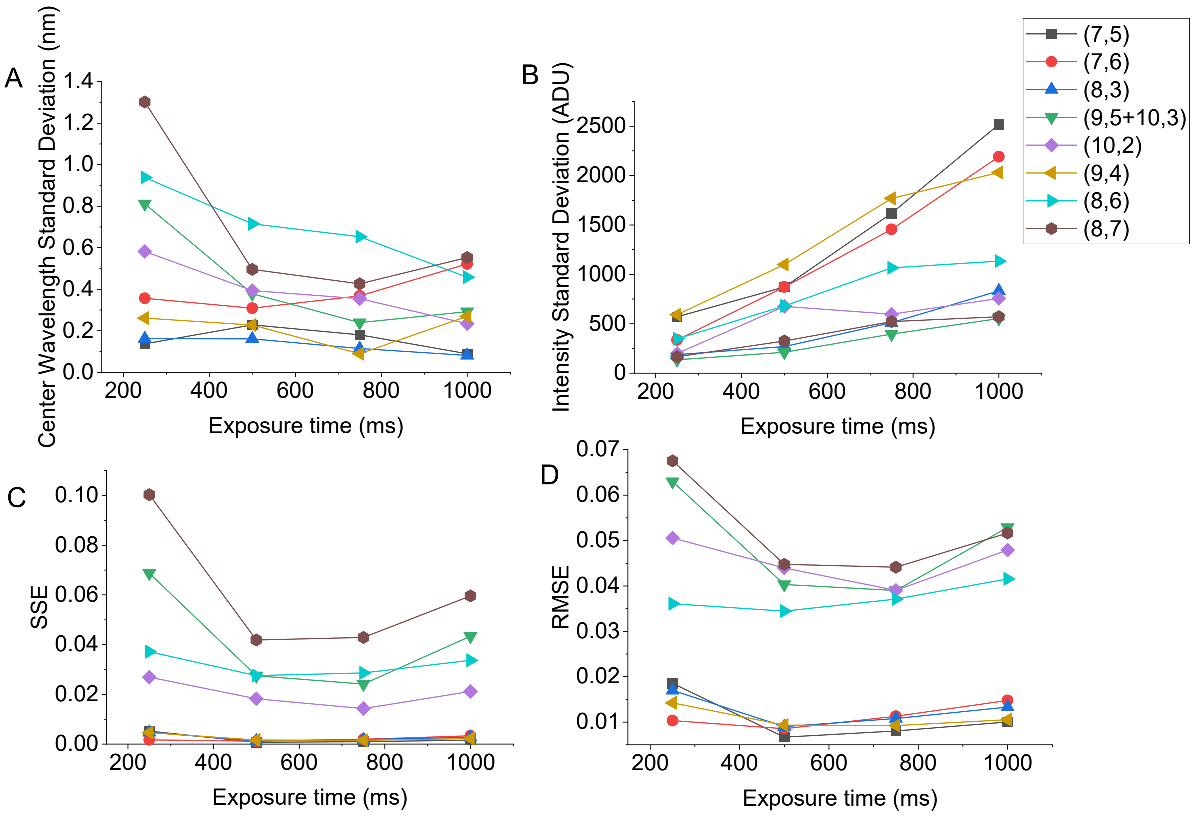


**Supplementary Figure 5**. **Statistical analysis of variations in exposure time of the spectral probe system.** A) Standard deviation in center wavelength of each chirality. B) Standard deviation in intensity of each chirality. C). standard square error (SSE) of each chirality. D) Root mean square error (RMSE) of each chirality. Laser power of 1000 mW was used for all time adjustments.

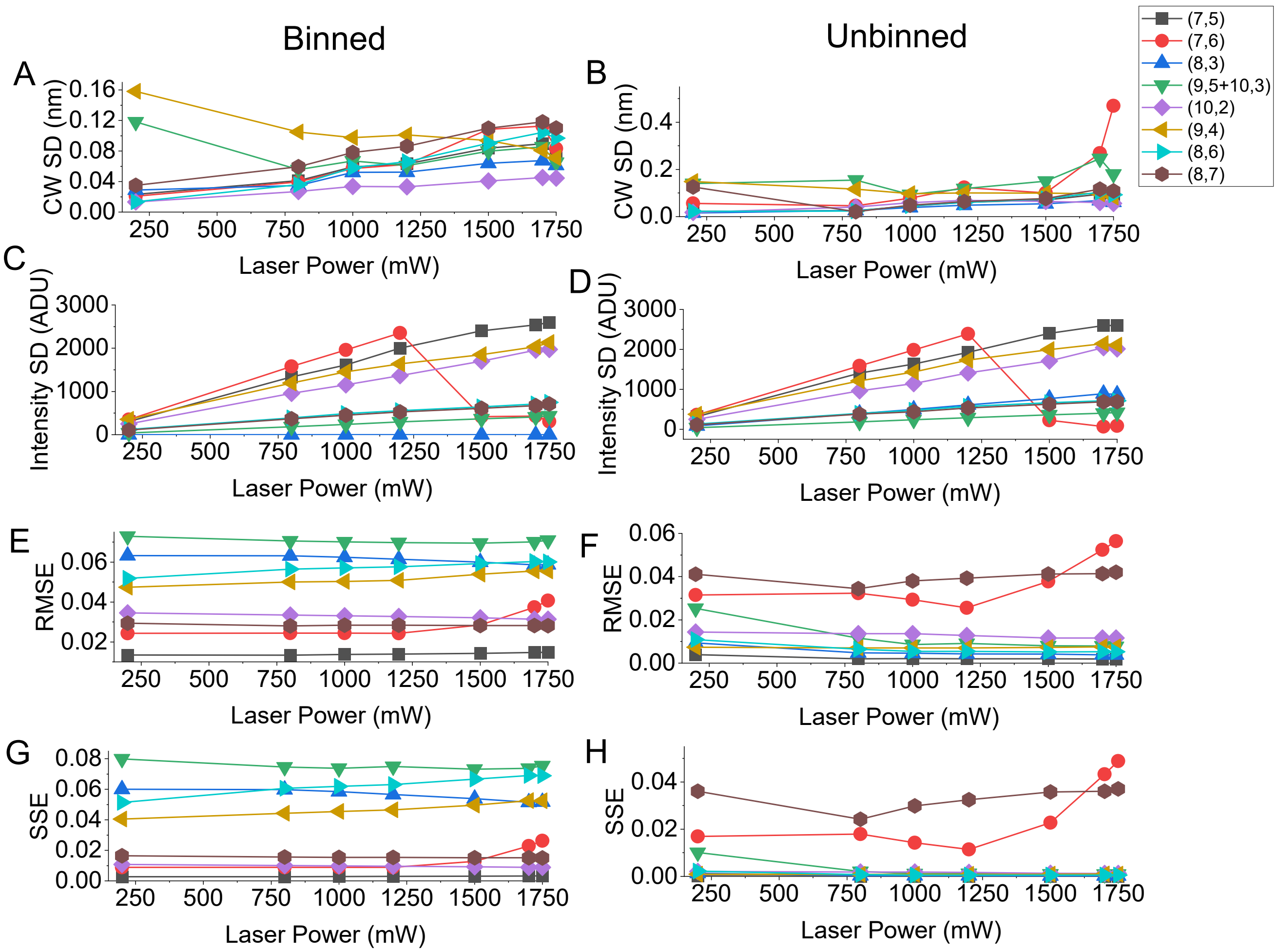


**Supplementary Figure 6. Comparison of analyses on binned and unbined data collected from varying laser power in the high thoughput system**. A) and B) standard deviation (n=12) of center wavelength values. C) and D) standard deviation (n=12) of intensity values. E) and F) root mean squared error values. G) and H) sum of squared error values
